# Heterozygous *KCNH2* variant phenotyping using Flp-In HEK293 and high-throughput automated patch clamp electrophysiology

**DOI:** 10.1101/2021.02.02.427891

**Authors:** Chai-Ann Ng, Jessica Farr, Paul Young, Monique J. Windley, Matthew D. Perry, Adam P Hill, Jamie I Vandenberg

**Affiliations:** Victor Chang Cardiac Research Institute, 405 Liverpool Street, Darlinghurst, NSW 2010, Australia; St Vincent’s Clinical School, UNSW Sydney, NSW, Australia

## Abstract

*KCNH2* is one of the 59 medically actionable genes recommended by the American College of Medical Genetics for reporting of incidental findings from clinical genomic sequencing. However, half of the reported *KCNH2* variants in the ClinVar database are classified as variants of uncertain significance. In the absence of strong clinical phenotypes, there is a need for functional phenotyping to help decipher the significance of variants identified incidentally. Here, we report detailed methods for assessing the molecular phenotype of any *KCNH2* missense variant. The key components of the assay include quick and cost-effective generation of a bicistronic vector to co-express WT and any *KCNH2* variant allele, generation of stable Flp-In HEK293 cell lines and high-throughput automated patch-clamp electrophysiology analysis of channel function. Stable cell lines take 3-4 weeks to produce and can be generated in bulk, which will then allow up to 30 variants to be phenotyped per week after 48 hours of channel expression. This high throughput functional genomics assay will enable a much more rapid assessment of the extent of loss of function of any *KCNH2* variant.

## Introduction

*KCNH2* encodes the human ether-à-go-go-related gene (hERG) potassium channel (Sanguinetti et al., 1995; Trudeau et al., 1995), which is responsible for the rapid component of the delayed rectifier potassium current, *I*_Kr_, a major contributor to cardiac repolarisation (Vandenberg et al., 2012). Loss-of-function variants in *KCNH2*, leading to reduced *I*_Kr_ current, result in prolongation of the cardiac action potential and cause congenital long QT syndrome type 2 (LQTS2) (Curran et al., 1995). Loss of function may be caused by reduced synthesis, misfold and reduced trafficking of channels to the cell membrane, altered gating or altered ion permeability of assembled channels (Delisle et al., 2004). Variants may also cause a dominant loss of function when co-expressed with WT alleles (Anderson et al., 2014).

Methods that are available to assess the *in-vitro* phenotype of *KCNH2* variants include the use of Western blot (Anderson et al., 2014; Perry et al., 2016), surface ELISA assay (Foo et al., 2019; Ng et al., 2020) or manual patch-clamp electrophysiology (Anderson et al., 2006; Ke et al., 2013). Patch-clamp assays are the gold standard for assessing function as they can assay levels of expression as well as altered gating and ion permeability all in the one assay. Until recently, patch clamp assays were limited by their low throughput (a few cells per day). This limitation, however, has been overcome by the introduction of high-throughput automated patch-clamp systems, which permit the analysis of the function for a large number of ion channel variants e.g. *KCNH2* (Ng et al., 2020), *KCNQ1* (Vanoye et al., 2018), *SCN5A* (Glazer et al., 2020), *KCNB1* (Kang et al., 2019) and CACNA1I (Heyne et al., 2020).

As hERG1a protein expresses as a tetrameric potassium ion channel potential dominant-negative effects will not be captured if phenotyping only the homozygous variant. To fill the unmet clinical need for the classification of clinically detected heterozygous *KCNH2* variants we need a high-throughput electrophysiology assay that can phenotype heterozygous *KCNH2* variants.

Here, we describe the methods for (1) designing the heterozygous expression vector; (2) generation of the stable Flp-In HEK293 cell lines in a high-throughput manner; (3) cell culture routine and cell preparation for use on automated patch-clamp systems. These protocols have been previously tested on a small number of heterozygous *KCNH2* variants (Ng et al., 2020) and has since been optimised into a high-throughput manner, including automated data analysis.

## Material and Methods

### Designing of DNA vector to express heterozygous *KCNH2* variant

To co-express variant and WT hERG1a protein we subcloned an internal ribosome entry site (IRES) into the pcDNA^™^5/FRT/TO vector (Thermofisher, cat. # V652020). The variant *KCNH2* cDNA was subcloned into the vector before the IRES and the WT *KCNH2* cDNA was subcloned after the IRES site. Previously, we showed using co-expression of hERG1a and hERG1b isoforms that placing the variant prior to the IRES and the WT after the IRES gave expression levels that most closely matched functionally equivalent expression of the two alleles (Ng et al., 2020). To maximise the efficiency of gene synthesis and subcloning of variants we introduced one restriction site (Nsil) into the variant *KCNH2* cDNA and silenced three restriction sites (*BstXl, BstEll* and *Sbfl*) in the WT *KCNH2* cDNA. Thus the variant *KCNH2* cDNA was divided into 5 regions, separated by *BstXl, BstEll, Nsil* and *Sbfl*, so that any heterozygous *KCNH2* variant vector can be generated from a synthesised gene block regardless of the location of the variant. We outsource this to GenScript Inc. (Pistcataway, NJ, USA) (see Fig. 1).

**Figure 1:**
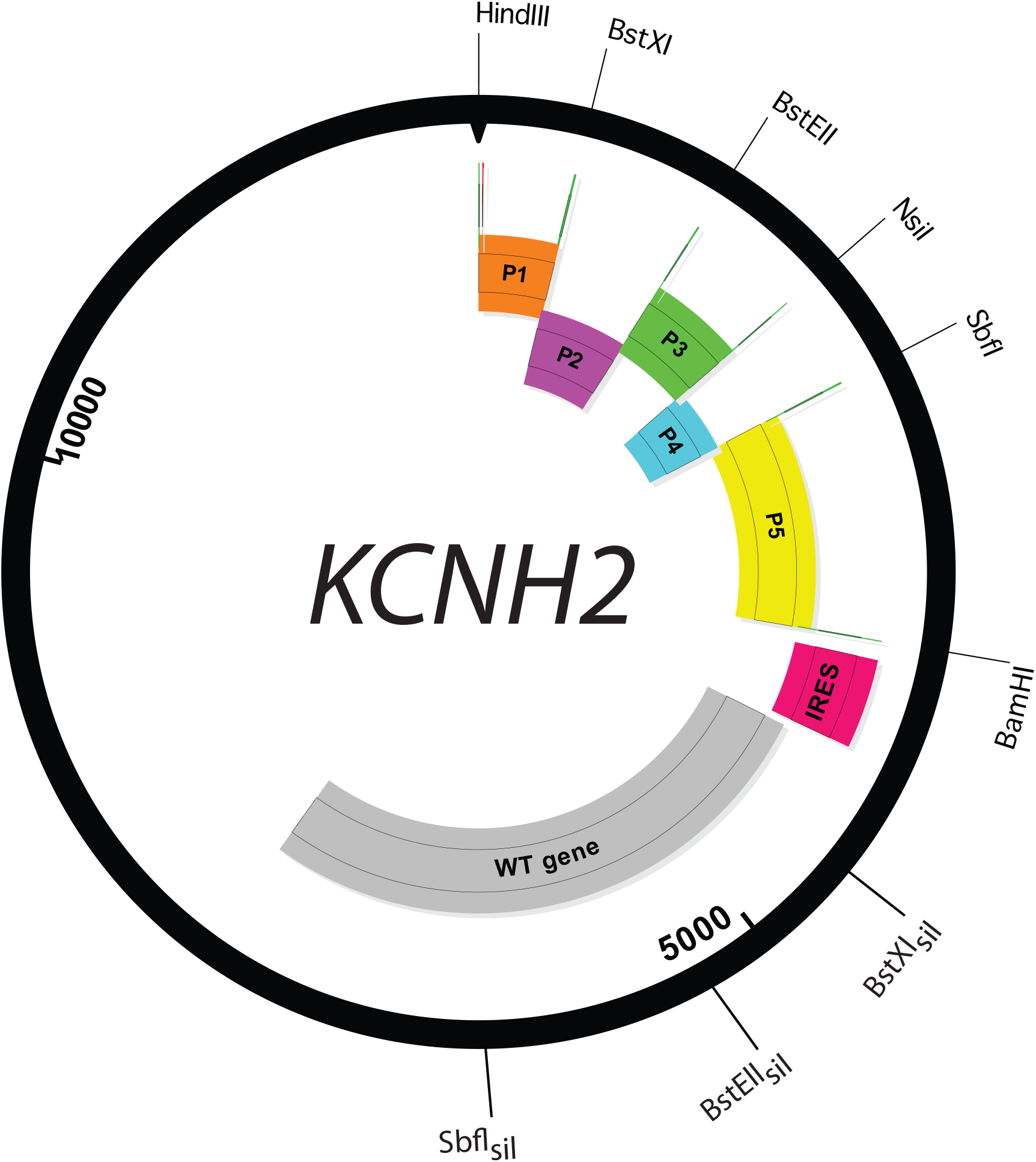
Design of a heterozygous *KCNH2* expression vector. *Nsil* was introduced into the *KCNH2* cDNA before the IRES while *BstXl, BstEll* and *Sbfl* were silenced in the *KCNH2* cDNA after the IRES. This allows the fragmentation of the *KCNH2* gene before the IRES into 5 sections to facilitate subcloning of variants.

### Generation of Flp-In™ T_rex_™ HEK293 *KCNH2* variant cell lines

The pipeline for generating a large number of Flp-In HEK293 *KCNH2* variants cell lines was optimised for establishment of 100 Flp-In HEK293 cell lines per round. The protocol is described as follow.

#### Day 0: Reviving Flp-In HEK293 parental cell (Thermofisher, cat. # R78007)

- A minimum of 40 million cells are required for 100 transfections.
- Thaw parental Flp-In HEK293 cells by incubating cryogenic vials in 37°C water bath for up to 3 minutes.
- Transfer cell suspension into a 15 mL centrifugate tube that contains 4 mL BZ-DMEM (DMEM (Thermofisher, cat. # 10566016) supplemented with 10 % FBS (Sigma-Aldrich, cat. # 12007C), 10 ug/mL Blasticidin (InvivoGen, cat. # ant-bl-1) and 100 ug/mL Zeocin (Thermofisher, cat. #R25005).
- Centrifuge cells at 200 g for 3 minutes. Aspirate old media and resuspend cells with BZ-DMEM, then dispense them into T75 flasks.
- Propagate cells to 80 % confluency.

#### Day 1: Prepare parental Flp-In HEK293 cells for transfection

- Passaging cells from T75 flasks.
  - *Aspirate media, wash with 5 mL DPBS (Thermofisher, cat. # 14190144) and aspirate. Digest cells with 1 mL TrypLE (Thermofisher, cat. # 12563011) and incubate flask at 37°C, 5 % CO*_*2*_ *for 4 minutes before adding 4 mL of BZ-DMEM to inactivate TrypLE*.
- Combine all cell suspensions into a T75 flask and count cells using Countess ll (Thermofisher, cat. # AMQAX1000). Transfer 40 millions cells into a fresh T75 flask and adjust the cell density to 200,000 cells per mL by adding fresh BZ-DMEM into the flask.
- Transfer 2 mL of cell suspension from the T75 flask into 101 wells (17 x 6-well plates), which will include 1 well for negative control.

#### Day 2: Transfection of heterozygous hERG vector DNA into parental Flp-In HEK293

- Prepare 100 wells (using 96-well plate format) that contain 520 ng of POG44 Flp-Recombinase Expression Vector (Thermofisher, cat. # V600520) and 2.3 uL of P3000 (Thermofisher, cat. # L3000015), add 60 ng of a different heterozygous *KCNH2* vector DNA into each well before adding 70 uL of Opti-MEM (Thermofisher, cat. # 31985070) into each well.
- In a separate set of 101 wells (1 additional well is for transfecting the negative control) (using 96-well plate format) add 2.3 uL of lipofectamine 3000 (Thermofisher, cat. # L3000015) and 70 uL of Opti-MEM into each well.
- Mix the DNA mixture and lipofectamine 3000 mixture by transferring the DNA mixture to the lipofectamine mixture and incubate at room temperature for at least 15 minutes.
- Replace the culture media of all the 6-well plates with 1.5 mL prewarmed Opti-MEM per well and add transfection reagent into each well.
- Return cells back into the 37°C, 5 % CO_2_ incubator for 5 hours before replacing the culture media with 2 mL B-DMEM (Thermofisher, cat. # 10566016) supplemented with 10 % FBS (Sigma-Aldrich, cat. # 12007C), 10 ug/mL Blasticidin (InvivoGen, cat. # ant-bl-1).

#### Day 3: Preparation of transfected Flp-In HEK293 for antibiotic selection process

- Passaging cells from each well into T25 flask
  - *Aspirate media, wash with 1 mL DPBS and aspirate. Digest cells with 0*.*15 mL TrypLE and incubate flasks at 37°C, 5 % CO*_*2*_ *for 4 minutes before adding 1 mL of B-DMEM to inactivate TrypLE*.
  - *Transfer all of the cell suspension from each well into a separate T25 flask that contains 4 mL of fresh B-DMEM*.

#### Day 4/6/8/10/12: Selection of stably integrated heterozygous *KCNH2* clones

- Replace the culture media in each T25 flask with BH-DMEM (DMEM supplemented with 10 % FBS, 10 ug/ml Blasticidin and 200 ug/mL Hygromycin) (Thermofisher, cat. # 10687010) at 2-day intervals.

#### Day 14: Propagation of stably integrated heterozygous *KCNH2* Flp-In HEK293 clones

- Passaging stably integrated clones into a new T25 flask.
  - *Aspirate media, wash with 2 mL DPBS and aspirate. Digest cells with 0*.*35 mL of a 1:1 diluted TrypLE with DPBS for 4 minutes at 37°C, 5 % CO*_*2*_ *incubator before adding 2 mL of BH-DMEM to inactivate TrypLE*.
  - *Transfer all of the cell suspension into a new T25 flask that contains 2*.*5 mL BH-DMEM*.

#### Day 18 to 25: Freezing down heterozygous *KCNH2* Flp-In HEK293

- Passaging cells when they reach 80 % confluency.
  - *Aspirate media, wash with 2 mL DPBS and aspirate. Digest cells with 0*.*35 mL TrypLE for 4 minutes at 37°C, 5 % CO*_*2*_ *incubator before adding 2 mL of BH-DMEM to inactivate TrypLE*.
  - *Transfer all of the cell suspension into a 15 mL tube and centrifuge at 200 g for 3 minutes*.
- Freezing cells
  - *Aspirate old media from the tube and resuspend cells with 2 mL cold BH-DMEM supplemented with 10 % sterile DMSO (Sigma-Aldrich, cat. # D2438)*.
  - *Dispense 1 mL per cryogenic vial and place them in an insulated container (Thermofisher, cat. # 5100-0001) for slow cooling. Place the insulated container in -80°C freezer and transfer the vials to liquid nitrogen vapour phase for long term storage the next day*.

### Cell culture routine of heterozygous *KCNH2* Flp-In HEK293 for automated patch clamp electrophysiology

The cell culture routine was established to allow the phenotyping of 10 variants and 2 controls (WT and negative controls) on a 384-well plate. This process can be up-scaled to 30 variants if desired.

#### Day 1: Reviving heterozygous *KCNH2* Flp-In HEK293 cells from cryogenic storage

- Thaw positive control (WT:WT), negative control (blank Flp-In HEK293) and 10 heterozygous variants by incubating these cryogenic vials in 37°C water bath for 3 minutes. This step is typically done on a Tuesday.
- Transfer each of the cell suspensions into a 15 mL centrifugation tube that contains 4 mL BH-DMEM and centrifuge at 200 g for 3 minutes.
- Aspirate old media and resuspend cells with 5 mL BH-DMEM and dispense them into individual T25 flasks.
- Propagate cells to near confluency, typically takes around 3 days.

#### Day 4/7/9/11: Recovery of heterozygous *KCNH2* Flp-In HEK293

- Passaging cells for 1 week to recover and synchronize cell growth
  - *Aspirate media, wash with 2 mL DPBS and aspirate. Digest cells with 0*.*35 mL TrypLE and incubate flasks at 37°C, 5 % CO*_*2*_ *for 4 minutes before adding 2 mL of BH-DMEM to inactivate TrypLE (negative control uses B-DMEM)*.
  - *Count cell using Countess ll and seed appropriate cell density into a fresh T25 flask that contains 4*.*5 mL of fresh BH-DMEM (negative control uses B-DMEM). For day 4 and day 11 passaging, seed 500k cells. For day 7 and day 9 passaging, seed 650k cells*.

#### Day 14: Seed heterozygous *KCNH2* Flp-In HEK293 for patch-clamp experiment

- Passaging 12 Flp-In HEK293 cell lines
  - *Aspirate media, wash with 2 mL DPBS and aspirate. Digest cells with 0*.*35 mL TrypLE and incubate at 37°C, 5 % CO*_*2*_ *for 4 minutes before adding 2 mL of BH-DMEM to inactivate TrypLE (negative control uses B-DMEM)*.
  - *Count cell using Countess ll and seed 1 million cells into a fresh T75 flask that contains 10 mL of fresh BH-DMEM (negative control uses B-DMEM)*.

#### Day 15: Induce the expression of heterozygous *KCNH2* Flp-In HEK293

- Induce expression by adding 200 ng/uL doxycycline (Sigma-Aldrich, cat. # D9891) into the media of each flask and propagate cells for 48-52 hours.

#### Day 17: Harvest cells for automated patch-clamp experiment

- Passaging 12 Flp-In HEK293 cell lines
  - *Aspirate media, wash with 5 mL DPBS and aspirate. Digest cells with 1 mL Accumax (Sigma-Aldrich, cat. # A7089) and incubate at 37°C, 5 % CO*_*2*_ *for 4 minutes*.
- Initial stage of cell recovery
  - *To recover cells following digest, add 5 mL of cold standard extracellular recording solution (in mM: NaCl 140, KCl 5, CaCl*_*2*_ *2, MgCl*_*2*_ *1, HEPES 10, D-Glucose 5; adjusted to pH 7*.*4 with NaOH) and incubate the flasks at 4°C for up to 15 minutes*.
- Single cell suspension
  - *Breaking cells into single cells by triturating the cell suspension 3-5 times and transfer the cell suspension into a 15 mL tube and centrifuge at 200 g for 3 minutes*.
  - *Aspirate the supernatant and resuspend cells with 5 mL of divalent-free extracellular recording solution (in mM: NaCl 140, KCl 5, HEPES 10, D-Glucose 5; adjusted to pH 7*.*4 with NaOH) in a 15 mL centrifugation tube, which will achieve a cell density between 0*.*5 to 1 million cells/mL depending on the confluency of cells. In our experience this range of cell densities does not affect the rate of cell catch on the SyncroPatch*.
- Final stage of cell recovery
  - *Rotate these tubes on a roller at 4°C for 30 minutes and transfer 3-4 mL of each cell suspension to a 12-well trough and rotate it at 300 rpm on a cell hotel that is maintained at 10°C on the SyncroPatch 384PE for up to 1 hour*.

### Operation of SyncroPatch 384PE automated patch-clamp

Operation of the SyncroPatch 384PE is according to the manufacturer’s instruction. Briefly, upon initiating the Biomek and PatchControl software, the robotic platform of Biomek is first reset by performing home axis and followed by executing the module “system flush” with water in PatchControl software after the tubes for the water and waste have been clamped. Place a box of tips and set up the internal solution (in mM: KF 110, KCl 10, NaCl 10, HEPES 10, EGTA 10; adjusted to pH 7.2 with KOH), 80 % Ethanol, divalent-free extracellular recording solution, high calcium extracellular recording solution (in mM: NaCl 140, KCl 5, CaCl_2_ 10, MgCl_2_ 1, HEPES 10, D-Glucose 5; adjusted to pH 7.4 with NaOH) and standard extracellular recording solution (Table 1). Enter the name for the 12 Flp-In HEK293 cell lines and complete the experiment definition and volumes for each of the solutions in the BioMek software. Place a 1-hole plate with medium resistance on the tray after it has been at room temperature for 15 minutes and start the experiment. The loading order of solutions/cells into the plate is as follow: divalent-free extracellular recording solution, internal solution, cells, high calcium extracellular recording solution, then two washing steps with standard extracellular recording solution to dilute the calcium back down to 2.75 mM. Cells were caught using –70 mbar whilst whole cell configuration is achieved by using a pressure of –250 mbar. Finally, voltage protocols are run. These can be configured for individual requirements. We typically record 1 s steady-state activation, onset of inactivation and 3s steady-state deactivation protocols (Ng et al., 2020). At the end of the recording, the plate is discarded and a technical replicate is performed with a new plate. The system is flushed with 80 % ethanol with a blank plate prior to system shut down.

### Voltage protocols and data analysis

Prior to analysing the current traces, all recordings were assessed using the following quality control parameters: seal resistance > 300 MΩ, capacitance between 5 and 50 pF, series resistance < 20 MΩ and leak corrected current measured at −120 mV leak step is within ± 40 pA of the baseline (Table 2). The QC assessment and data analysis were performed using custom written python scripts (see https://git.victorchang.edu.au/projects/SADA/repos/syncropatch_automated_analysis/browse; a sample data for WT is also available at https://doi.org/10.26190/7tqy-gc11). In addition, if the number of wells that pass QC for analysis of gating parameters is less than 10 % of the recordings that have passed the plate recording QC parameters then these variants are classified as not analysed (N/A) for the relevant gating parameters in the result.

#### Peak current density

Cells were depolarised to +40 mV for 1 s then hyperpolarized to −120 mV. Peak current density was measured as the peak amplitude of the tail current (see Fig. 4A, asterisks indicate where the peak tail current amplitudes were measured). Membrane capacitance was used to normalise the current density for comparison between recordings.

**Figure 2:**
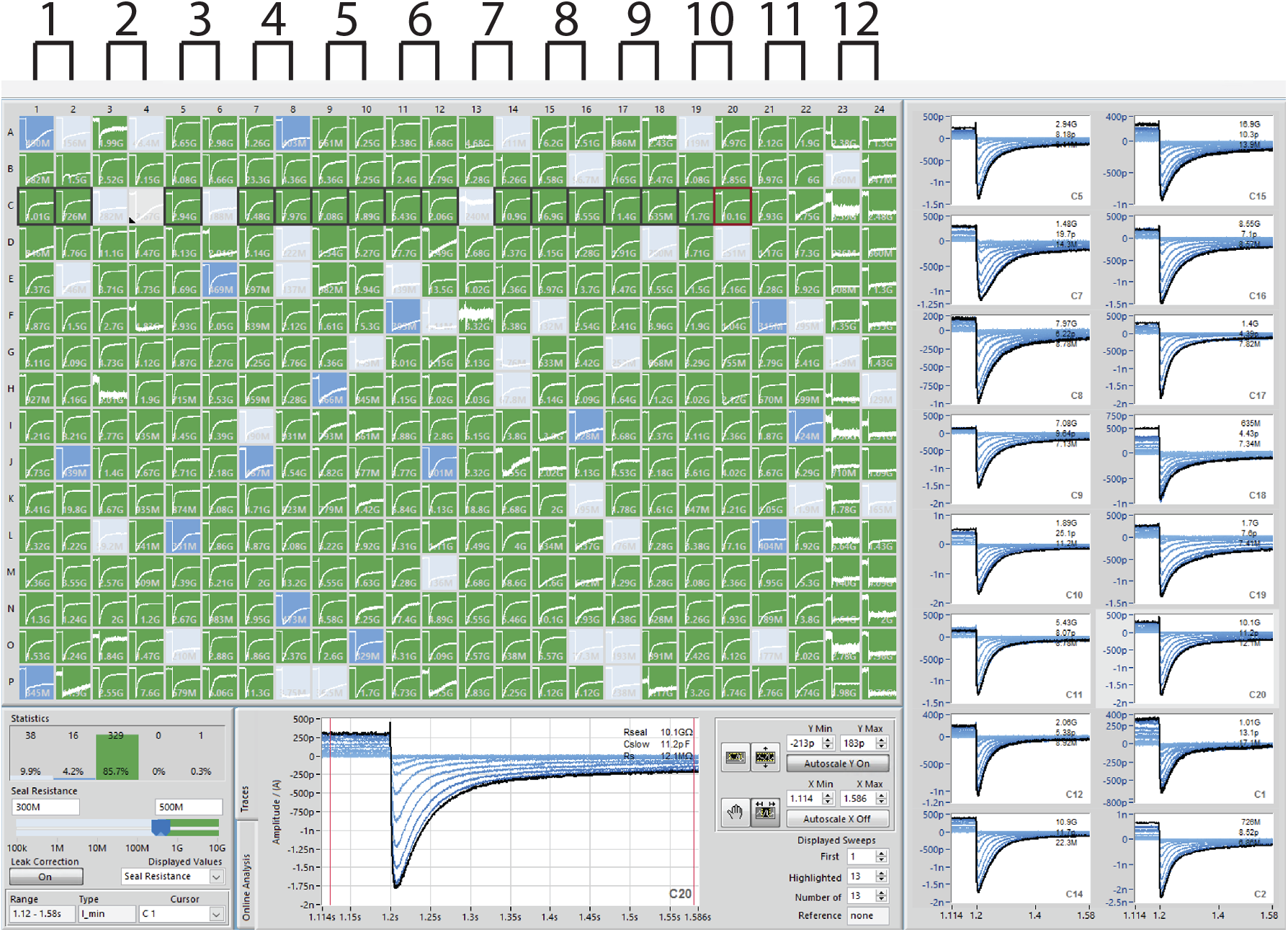
Overview of PatchControl software. Example of −120 mV tail currents for unmodified (column 1 to 6) and modified (column 7 to 11) WT hERG channels, as well as, a negative control (column 12). In this example, out of a total of 384 wells, 86 % of wells have seal resistance > 500 MΩ (green), 4 % have a seal resistance in the range 300 -500 MΩ and 10 % of wells have seal resistance <300 MΩ (grey).

**Figure 3:**
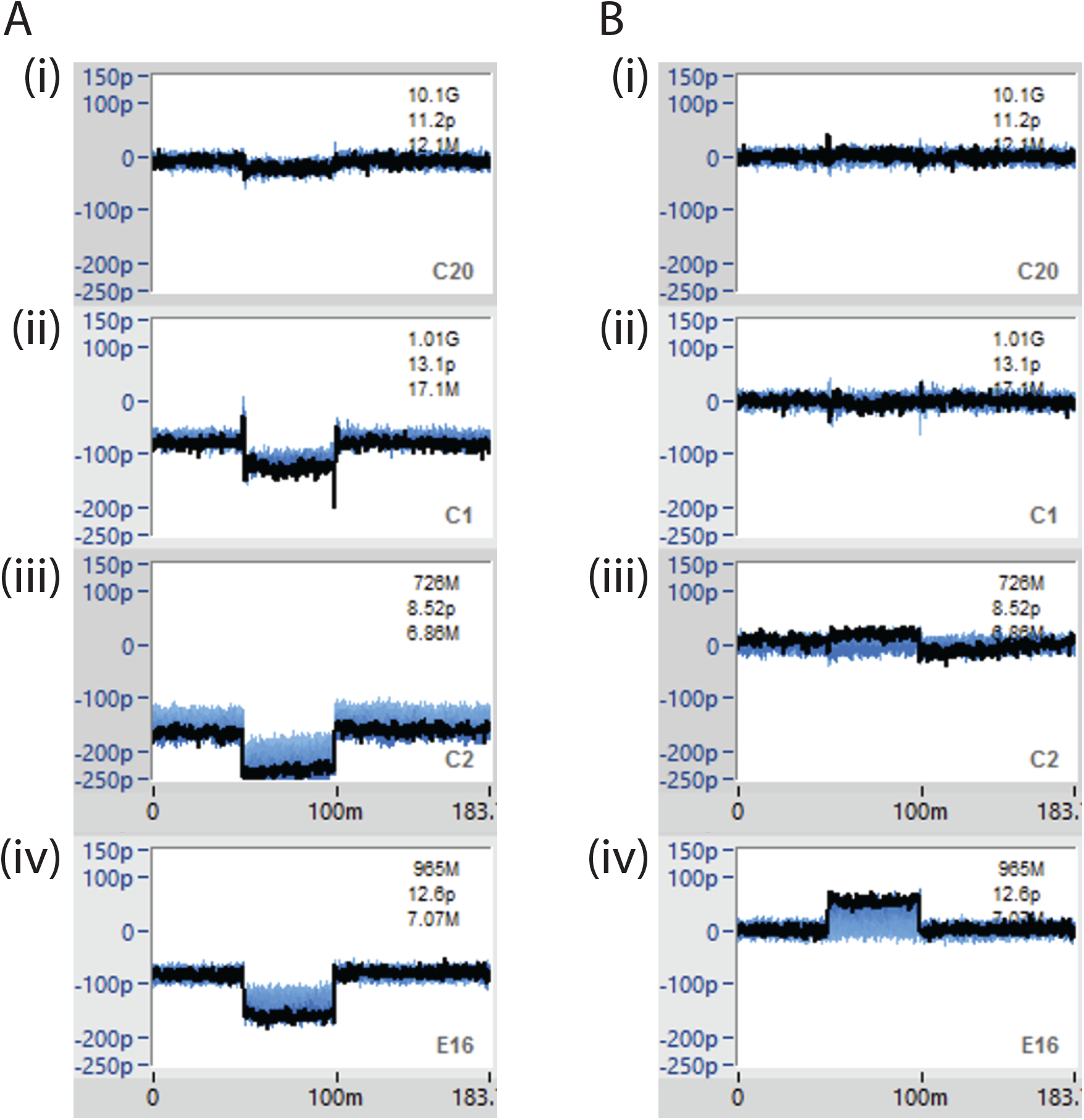
Examples of leak current correction. (A) uncorrected and (B) corrected current traces. Only corrected current traces that have median current value that are less than ±40 pA from the baseline for the region between 60-80 ms in each protocol are accepted for further analysis. Panels (i)-(iii) pass this QC criterion whereas panel (iv) would fail.

**Figure 4:**
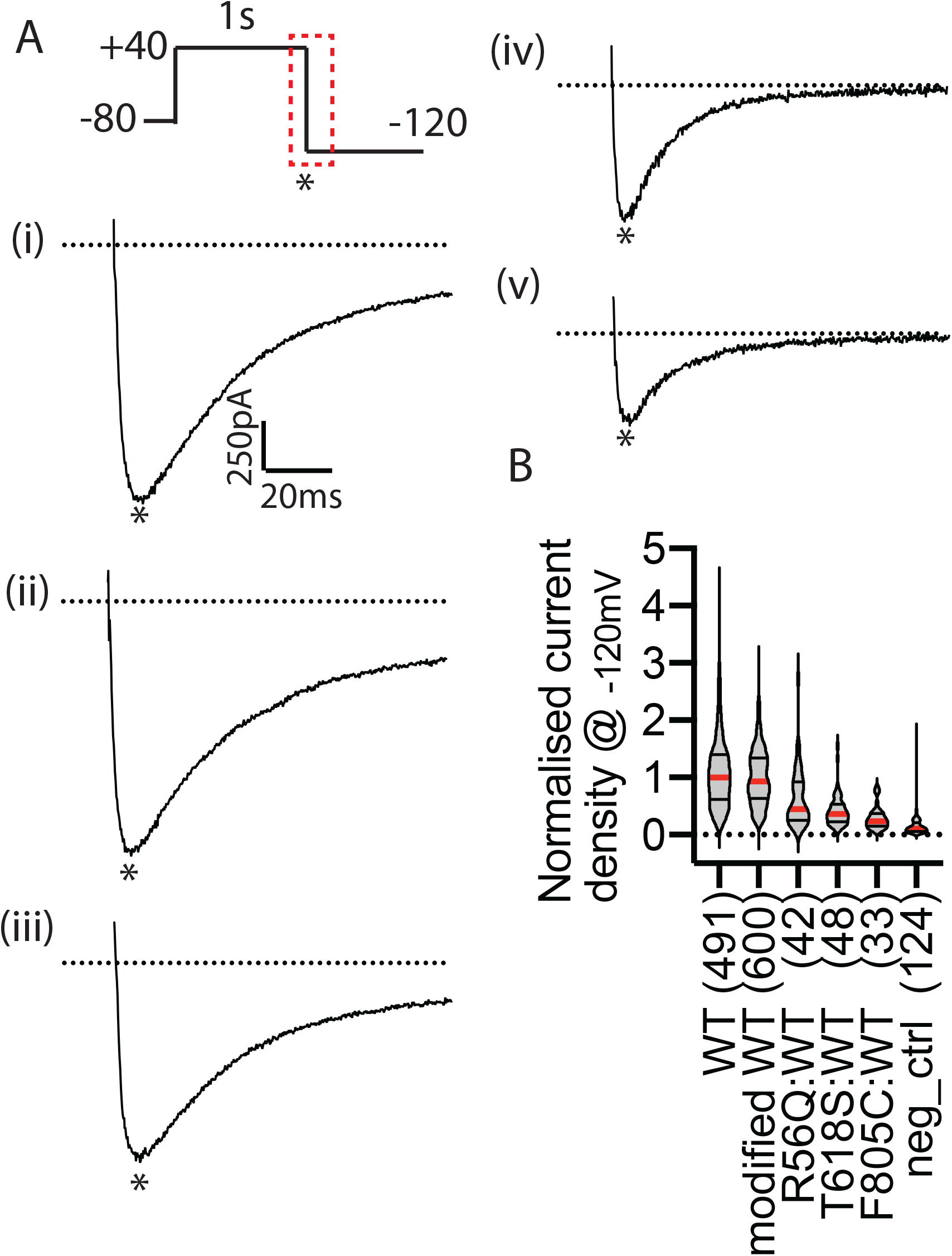
Current density measurements. (A) Examples of −120 mV tail current traces for (i) unmodified WT, (ii) modified WT, (iii) R56Q:WT, (iv) T618S:WT and (v) F805C:WT. Voltage protocol is shown as inset on panel (ii) and the dashed lines indicate region of the protocol corresponding to the current traces shown. The peak current is indicated by asterisk. (B) Summary of current densities for all constructs shown as violin plot with median indicated by the red horizontal bar with upper and lower quartiles shown as black bars.

#### *V*_0.5_ of activation

The voltage dependence of activation was measured using a 1 s isochronal activation protocol; cells were depolarised to potentials from –50 mV to +70 mV in 10 mV increments for 1 s before stepping to –120 mV to record tail currents (see Fig. 5A, the tail currents correspond to the dashed box highlighted in the voltage protocol). To determine the mid-point of the voltage dependence of activation (V_0.5_), the −120 mV peak tail currents recorded after each depolarizing voltage step, I, were fit with a Boltzmann distribution using Equation 1:

**Figure 5:**
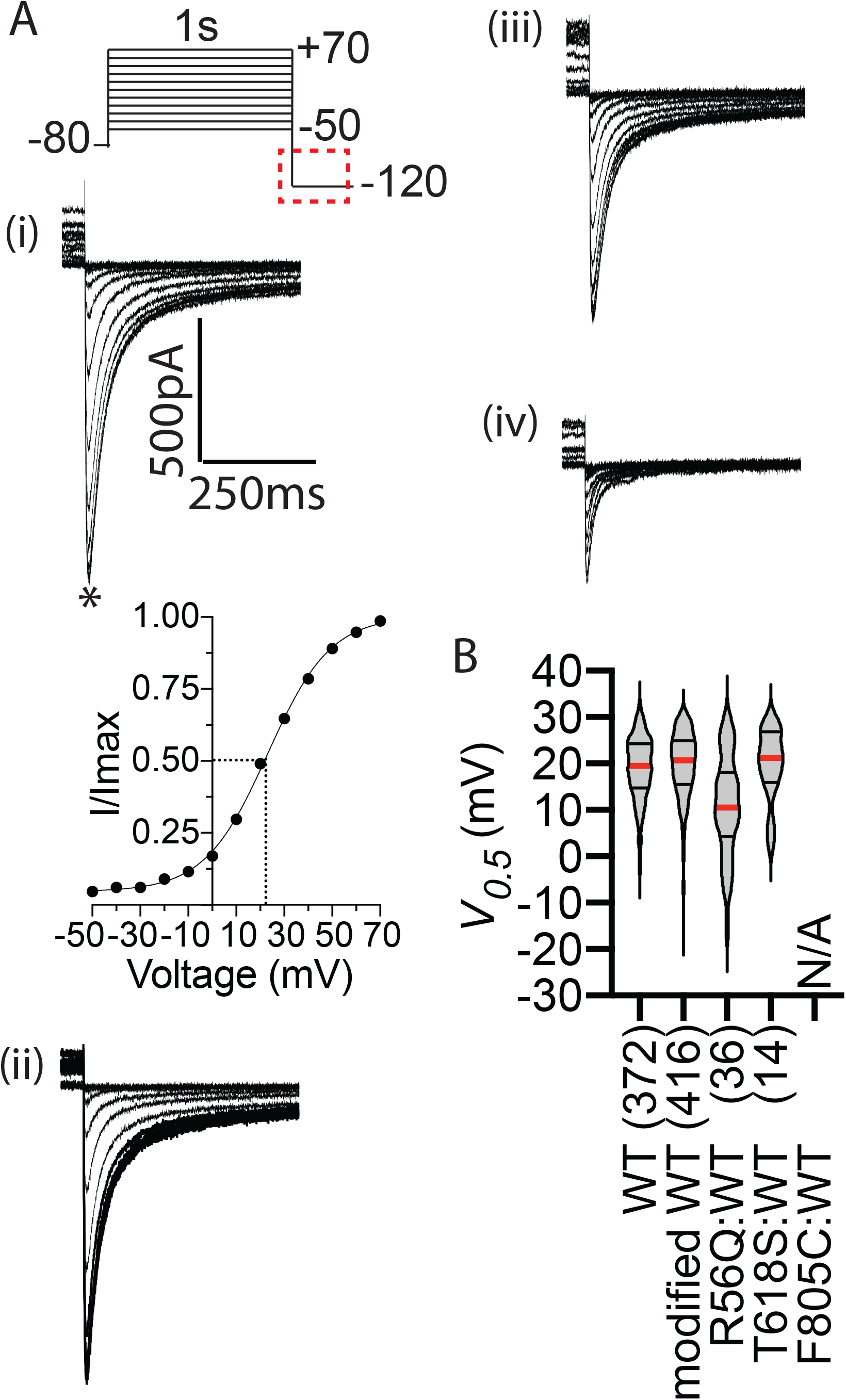
Activation phenotyping. (A) Example of −120 mV tail current traces recorded for (i) unmodified and (ii) modified WT *KCNH2*, (iii) R56Q:WT and (iv) T618S:WT following 1 s of depolarisation from the holding potential to voltages in the range −50 to +70 mV. Dashed box on the voltage protocol highlights the portion of the protocol depicted in the current traces. Example of Boltzmann fit to the normalised peak current indicated by asterisk in (i) to show how the V_0.5_ of activation is calculated. (B) Summary of V_0.5_ values for steady-state activation derived from fitting a Boltzmann function to the peak tail current amplitudes (see Methods for details). The median values are indicated by the red horizontal bar with upper and lower quartiles shown as black bars. N/A indicates when the n number is less than 10 % of recordings that pass the QC.

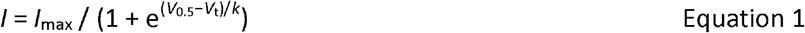

where: *V*_0.5_ is the half-maximum activation voltage, *V*_t_ is the test potential, *k* is the slope factor and *I*_max_ is the maximum tail current (see Fig. 5Ai).

#### Channel deactivation

The rate of hERG channel deactivation was measured by depolarising cells to +40 mV for 1 s before repolarising cells to potentials in the range +20 to −150 mV in 10 mV decrements for 3 s (see Fig. 6A, the tail currents correspond to the dashed box highlighted in the voltage protocol). The tail current represents the recovery from inactivation followed by channel deactivation. To determine the time constant of deactivation, the decaying portion of the tail current recordings were fit with a double exponential function (red line). The overall time constant of deactivation was calculated as a weighted sum of the two components (i.e. fast and slow components of deactivation) using Equation 2:

**Figure 6:**
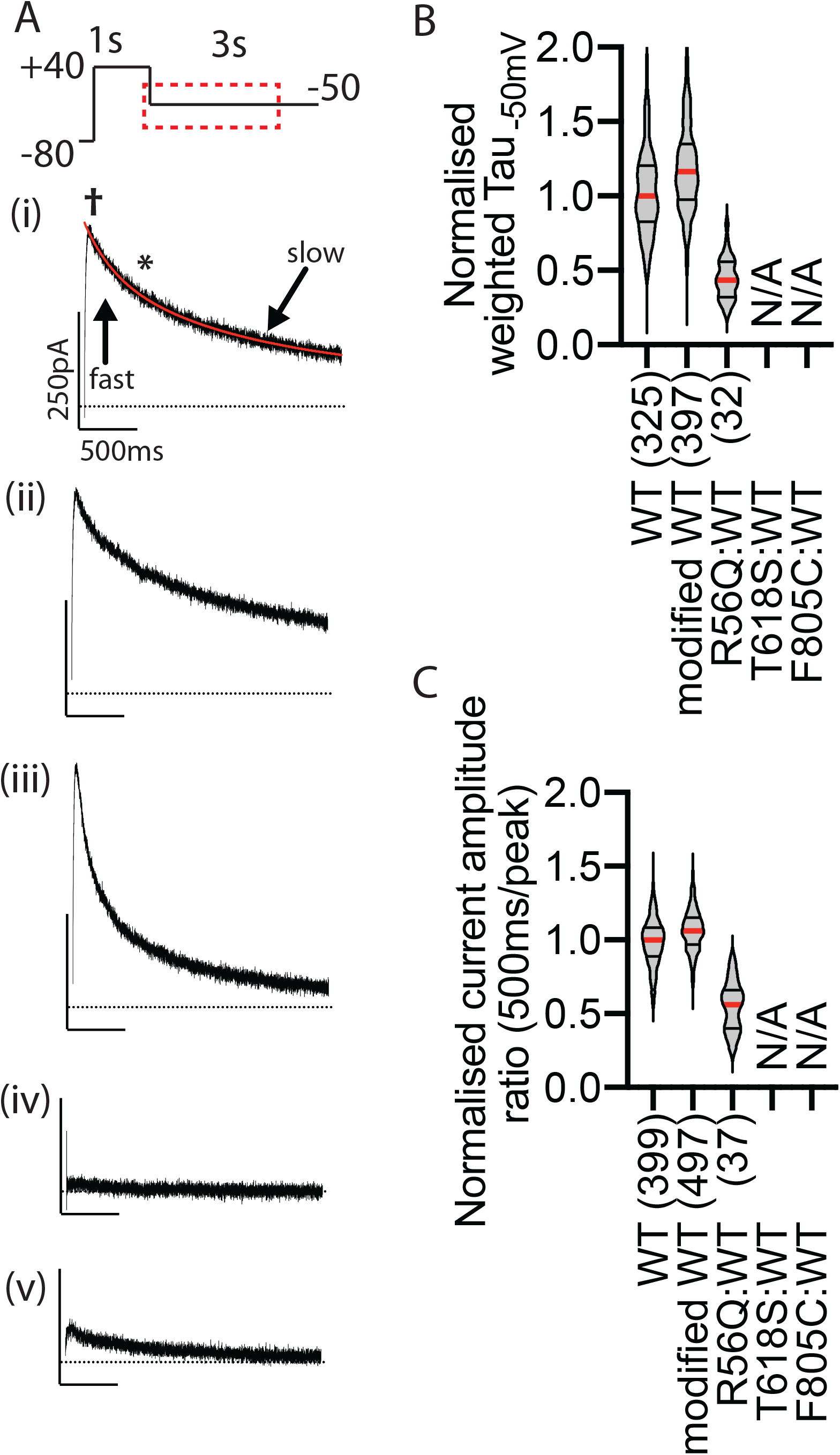
Deactivation phenotyping. (A) Example −50 mV tail current traces (corresponding to dashed line within the voltage protocol) for (i) unmodified and (ii) modified WT *KCNH2*, (iii) R56Q:WT, (iv) T618S:WT and (v) F805C:WT. The double exponential fitting of the deactivation decay is shown as red in (i) with arrows indicating the fast and slow components. (B) Summary of weighted time constant for channel deactivation (see methods for details). The median values are indicated by the red horizontal bar with upper and lower quartiles shown as black bars. It was only possible to analyse those constructs where the peak tail current were greater than 100 pA, hence T618S:WT is excluded and F805C:WT has limited numbers. (C) Summary of normalised current amplitude ratio where the current amplitude at 500 ms after the peak current (* in panel i) is divided by the peak current (†). N/A indicates when the n number is less than 10 % of recordings that pass the QC.

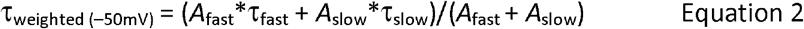

where *A* is the current amplitude and τ is the time constant.

We also directly measured the extent of channel deactivation at 500 ms, by measuring the peak current at −50 mV (indicated by † in Fig. 6Ai) and 500 ms later (indicated by * in Fig. 6Ai), which is reported as the current decay ratio at 500 ms (see Fig. 6C).

#### Channel inactivation

The rate of onset of inactivation was measured using a triple pulse protocol where cells were depolarised to +40 mV for 1 s to fully activate and then inactivate the channels before stepping to −110 mV for 10 ms to allow channels to recover from inactivation into the open state. Membrane potential was then depolarised to voltages in the range +60 to −20 mV (in 10 mV decrements) to inactivate channels (see Fig. 7A, the tail currents correspond to the red line highlighted in the voltage protocol). To determine the time constant for the onset of inactivation, we fitted a single exponential to the inactivating current trace (red line). To minimise the impact of poor voltage clamp control during large outward current traces we only fitted the exponential function to the current trace after it had fallen to <500 pA. A second estimate of the effects of variants on inactivation were obtained by measuring the ratio of the peak tail current amplitudes at −50 mV (indicated by * in Fig. 7C) and −120 mV (indicated by † in Fig. 7C) recorded from the deactivation protocol (see above).

**Figure 7:**
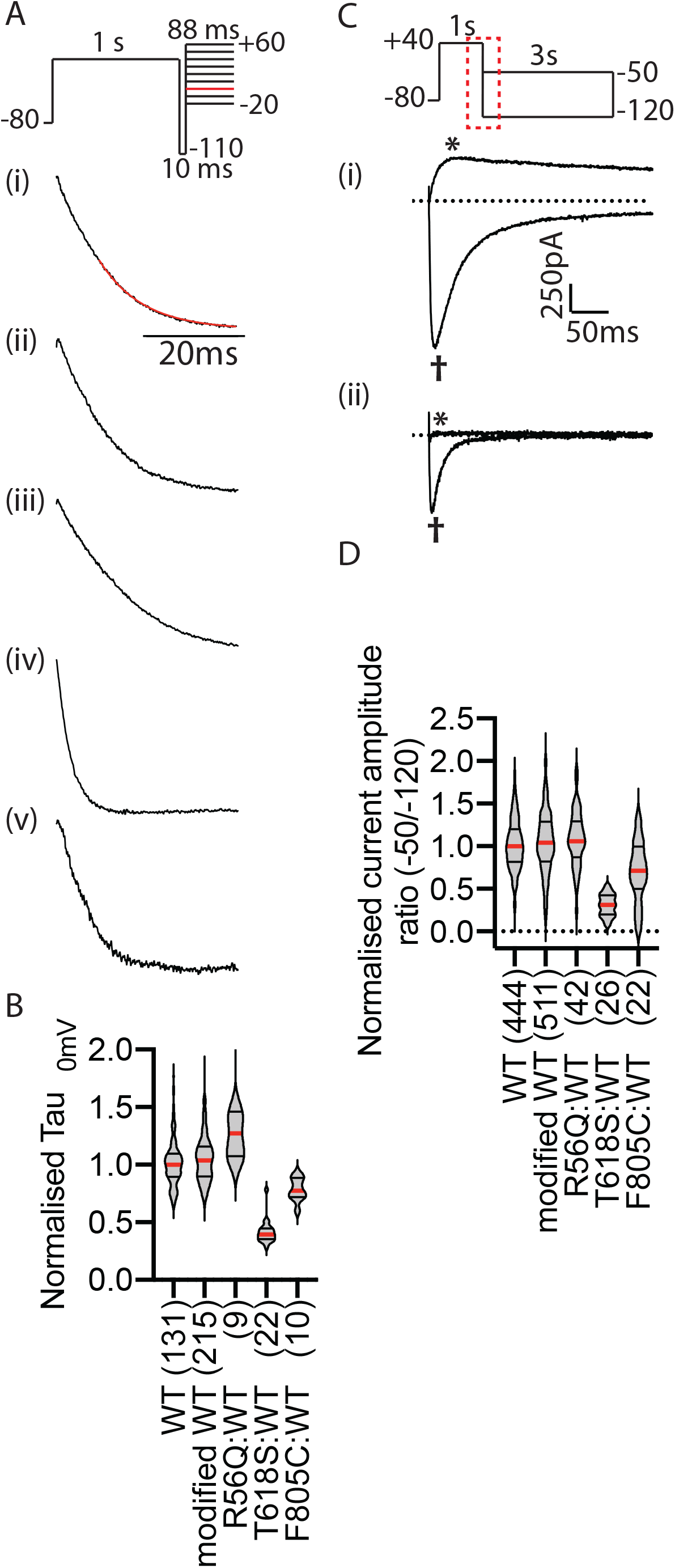
Inactivation phenotyping. (A) Normalised current traces for the onset of inactivation. The dashed box within the voltage protocol highlights the region shown in the current traces. (i) unmodified and (ii) modified WT *KCNH2*, (iii) R56Q:WT, (iv) T618S:WT and (v) F805C:WT. The single exponential fitting of current decay from 500 pA was shown as red in (i). (B) Summary of tau constants for the onset of inactivation at 0 mV. The median values are indicated by the red horizontal bar with upper and lower quartiles shown as black bars. (C) Example tail currents recorded at −50 mV (outward currents) and −120 mV (inward current) for modified WT *KCNH2* (black), and T618S:WT (red). (D) Summary of the normalised current amplitude ratio (i.e. */† in C) for all constructs. The smaller value for the T618S:WT constructs is indicative of enhanced inactivation of this variant. The median values are indicated by the red horizontal bar with upper and lower quartiles shown as black bars.

## Results and Discussion

To enable cost-effective gene fragment synthesis, Nsil was introduced into the variant *KCNH2* cDNA and *BstXl, BstEll* and *Sbfl* within the WT *KCNH2* cDNA were silenced. Together with the 5’ *Hindlll* and 3’ *BamHl*, gene synthesis of any fragments (either P1, P2, P3, P4 or P5) that harbour specific *KCNH2* variants can be synthesised and subcloned into the heterozygous *KCNH2* vector (Fig. 1). The heterozygous *KCNH2* vector can then be used to generate isogenic heterozygous *KCNH2* Flp-In HEK293 cell lines for functional phenotyping (Ng et al., 2020).

As the phenotyping patch-clamp experiment is performed using a high-throughput automated patch-clamp system (SyncroPatch 384PE), we have optimised the high throughput generation of Flp-In HEK293 cell lines as well. A total of up to 100 Flp-In HEK293 cell lines can be generated per round (see methods) with very high efficiency when individual transfection reagents are used (i.e. lipofectamine 3000 and P3000 reagent were prepared individually for each transfection using multichannel pipettes in a 96-well format). Typically, stable isogenic heterozygous *KCNH2* Flp-In HEK293 clones will be ready for cryo preservation 3 weeks post transfection. Generation of stably integrated heterozygous *KCNH2* doxycycline inducible Flp-In HEK293 cell lines enabled us to obtain a very high number of cells that are expressing the hERG potassium channel. In this study, of the 2112 wells seeded with *KCNH2* Flp-In HEK293 cells, 1908 wells had seal resistance greater than 300 MΩ (∼90 % of wells). Out of these wells, 1841 wells (96.5 %) have “hooked” tail currents. This compares very favourably with previous analyses of transient transfection which achieved ∼80 % transfection efficiency as detected by flow cytometry (Vanoye et al., 2018).

To phenotype heterozygous *KCNH2* Flp-In HEK293 cell lines we used a SyncroPatch 384PE automated patch-clamp electrophysiology system (Nanion Technologies, Munich, Germany). We found that the medium resistance, single hole chips, were suitable for performing the patch-clamp experiment on Flp-In HEK293 cells. It is worth noting that the incubation of these Flp-In HEK293 cells with cold standard extracellular recording solution immediately after the treatment with accumax (see methods) is essential to achieve a good seal resistance; in our experience this enables >80 % with seal resistances greater than 500 MΩ (green panels in Fig. 2). However, it is worth noting that this cold incubation step is likely to be cell type and cell line dependent.

The SyncroPatch assay was designed to phenotype 10 variants in parallel (column 2 to 11 in Fig. 2) together with a positive (column 1; WT) and a negative control (column 12; blank Flp-In HEK293), i.e., the assay is designed to produce equal number of patch-clamp experiments for WT, variants and negative control per plate. Each experiment is also run in duplicate to achieve a total of 64 patch-clamp experiments per variant.

The leak current can be corrected by the PatchControl software to produce leak-corrected current traces but to prevent over or under correction, a 50 ms step at −120 mV was implemented at the beginning for all protocols. Recordings were excluded from analysis when the leak corrected current at this –120 mV is greater than ± 40 pA (Fig. 3). Most of the recordings corrected by the PatchControl software is good (panel i and ii) or acceptable (panel iii). Overall, in the WT data presented in this study, < 1 % of wells were excluded due to failing the ± 40 pA threshold criterion (see Table 2).

As the restriction sites of the heterozygous *KCNH2* vector were modified to WT_Nsil_:WT_(BstXl/BstEll/Sbfl)-silenced_ to allow affordable gene fragment synthesis and subcloning of variants, we first compared our modified WT construct with an unmodified WT:WT hERG1a, which shows the modification of the restriction sites has not affected the channel expression level (Fig. 4B). We also included three other *KCNH2* variants with gating or trafficking defects (R56Q – accelerated deactivation, T618S – enhanced inactivation and F805C – trafficking deficient) to illustrate how the molecular phenotypes of different *KCNH2* variants can be assessed using the automated patch-clamp phenotyping assay.

We first measured the tail current amplitude at –120 mV after the channel was depolarised at +40 mV for 1 s (Fig. 4A). In this example, F805C (panel v) is a known trafficking deficient variant (Ficker et al., 2002) and the analysis of tail current amplitude can unambiguously identify its loss-of-function phenotype (Fig. 4B). In addition to the loss-of-function phenotype as a result of having reduced hERG expression, abnormal hERG channel gating can also reduce hERG function. Steady-state activation is the transition of the channel from closed to open conformation and the V_0.5_ of activation can be measured by fitting the tail current amplitudes using Boltzmann function (Fig. 5A). R56Q has a modest shift in the V_0.5_ of activation (–10 mV) whilst the other variants tested in this study had WT-like V_0.5_ values (Fig. 5B).

For those variants that are capable of passing current at –50 mV (Fig. 6A, panels i-iii), some of them might accelerate channel deactivation (i.e. transition from open to closed conformation more rapidly) and cause loss-of-function phenotype (panel iii). The decay of the current can be fitted using a double exponential function to derive the weighted time constant of channel deactivation. For example, R56Q deactivates twice as fast compared to WT channels (Fig. 6B). The weighted time constant include both the fast and slow components of channel deactivation. In addition, the current amplitude ratio is taken at 500 ms from the peak, which predominantly reflects the fast component (Fig. 6C) and has the capability to detect those variants that have slower channel deactivation in the physiological time range. Identifying fast deactivating variants is important as these variants can reduce protective *I*_Kr_ current that is critical for suppressing premature depolarisations or ectopic beats (Lu et al., 2001).

For the two variants that are incapable of passing current at –50 mV (see Fig. 6), F805C is mostly trafficking defective whilst T618S has an enhanced inactivation (Fig. 7A, panel iv and Fig. 7B). This enhanced inactivation causes loss-of-function as the channel does not pass current at –50 mV (Fig. 7C, panel ii). This loss-of-function phenotype can also be quantified by taking the ratio between the –50 and –120 mV tail current amplitudes (Fig. 7C). A smaller value than control is an indicative of loss-of-function phenotype due to enhanced inactivation phenotype (Fig. 7D).

From the example data we have presented in this study it is also possible to estimate the power of the assay to detect a 25 %, 50 % or 75 % loss-of-function in current density. For 80 % power with an alpha of 0.05, and effect sizes of 0.47, 0.94 and 1.41, respectively (determined based on the standard deviation for the WT data) we would require n = 35, 9, 4 to detect a 25, 50 % or 75 % loss-of-function.

In conclusion, we have developed an efficient workflow that allows high-throughput functional phenotyping of any *KCNH2* variant by integrating specially design cloning strategy, generation of heterozygous *KCNH2* Flp-In HEK293 cell lines and high-throughput automated patch-clamp electrophysiology (Fig. 8). This phenotyping assay has the potential to assist in the classification of *KCNH2* variants, which will hopefully improve the diagnosis and management of LQTS2 patients and more importantly reduce the number of variants that are classified as VUS. However, before this can be considered it will be important to establish the robustness of this assay in predicting the loss of function of variants for which there is already strong clinical data to indicate whether the variants are pathogenic or benign as per the recommendations proposed by Brnich et al. (2019).

**Figure 8:**
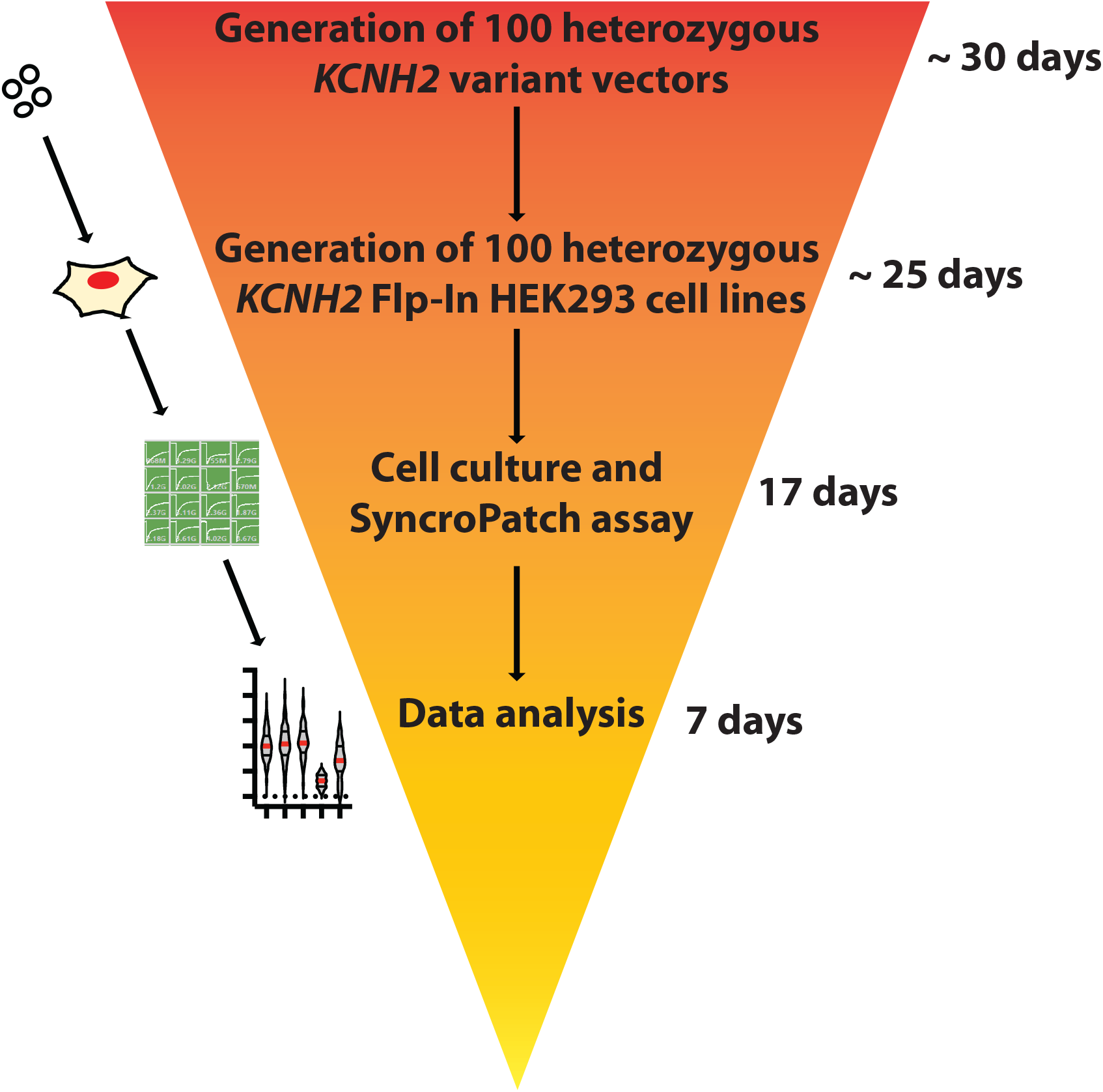
Typical timeline for *KCNH2* phenotyping assay. Generation of heterozygous *KCNH2* vectors are typically done in a batch of 100. We outsource this process to GenScript Inc (Pistcataway, NJ, USA), which has a turnaround of ∼30 days. Generation of heterozygous *KCNH2* Flp-In HEK293 cell lines will take around 25 days. Cell culture and the SyncroPatch assay takes 17 days which is done in batches of 10 variants staggered at weekly intervals. Data analysis and curation takes ∼7 days per batch of variants. In total, it will take less than 3 months from generating the variant vector to the final data analysis.

## Acknowledgements

We would like to thank Jeffrey McArthur and acknowledge the Victor Chang Cardiac Research Institute Innovation Centre, funded by the New South Wales Government, Australia. We also thank Soren Wohlthat, Hank Levsen and Richard Cooke for help with development of analysis scripts.

