## Supplementary material for "Heterozygous *KCNH2* variant phenotyping using Flp-In HEK293 and high-throughput automated patch clamp electrophysiology": Table 1 and 2

**Table 1: Recording solutions**

| mM | NaCl | KCl | CaCl_2_ | MgCl_2_ | HEPES | Glucose | KF | EGTA |
| --- | --- | --- | --- | --- | --- | --- | --- | --- |
| Internal  (pH 7.2 with KOH) | 10 | 10 |  |  | 10 |  | 110 | 10 |
| Divalent-free extracellular (pH 7.4 with NaOH) | 140 | 5 |  |  | 10 | 5 |  |  |
| High calcium extracellular (pH 7.4 with NaOH) | 140 | 5 | 10 | 1 | 10 | 5 |  |  |
| Standard extracellular (pH 7.4 with NaOH) | 140 | 5 | 2 | 1 | 10 | 5 |  |  |

**Table 2: Quality control parameters for the 6 WT plates (including negative control)**

| Seal resistance | >300 MΩ | 2083/2304 wells |
| --- | --- | --- |
| Capacitance | 5-50 pF | 1381/2083 wells |
| Series resistance | <20 MΩ | 1206/1381 wells |
| Leak corrected current | ± 40 pA of the baseline | 1201/1206 wells |
| % of wells passing all QC |  | 1201/2304 (52.1 %) |
